# A flow cytometry protocol for accurate and precise measurement of plant genome size using frozen material

**DOI:** 10.1101/2024.02.14.580322

**Authors:** Abhishek Soni, Lena Constantin, Agnelo Furtado, Robert J Henry

**Affiliations:** Queensland Alliance for Agriculture and Food Innovation, The University of Queensland St Lucia Qld 4072 Australia; ARC Centre of Excellence for Plant Success in Nature and Agriculture, The University of Queensland St Lucia Qld 4072 Australia

**Keywords:** Flow cytometry, genome size, nuclear isolation, frozen plant material, debris compensation, histogram modelling

## Abstract

Flow cytometry is a technique widely applied to infer the ploidy and genome size of plant nuclei. The conventional approach of sample preparation, reliant on fresh plant material to release intact nuclei, requires protocol optimisation for application to many species. The approach often results in poor yields of nuclei, impeding the accurate measurement of genome size and confines the optimal resource allocation and efficiency in genome sequencing which relies on genome size estimation. Here, we present a novel method using frozen plant material that facilitates the release of intact nuclei for genome size estimation. Genome estimates from frozen material are similar to those from fresh material. Accurate and precise estimates can be made by complementing the fluorescence of frozen nuclei with histogram modelling and debris compensation algorithms. This method of nuclei isolation from frozen plant material for flow cytometry-based genome size estimations has special value in estimating the genome size of samples collected and frozen for use in plant genome sequencing. Plant material can be conveniently stored, resampled, and used for DNA or RNA extractions.

**Highlight:** Frozen leaf material can be used to isolate nuclei for the accurate estimation of genome size The method proved suitable for difficult samples and did not require specific optimization. The method was especially useful where plant material could not be immediately processed through flow cytometry and allowed the same sample to be used for genomes size estimation and genome sequencing.

## Introduction

Flow cytometry (FCM) is widely used to estimate genome size (GS) and ploidy in plants. The method involves the preparation of a suspension of intact nuclei, labelling the nuclei with a fluorochrome that binds to nucleotides, and measuring the fluorescence intensity of each nuclei (Doležel *et al*., 2007; Galbraith *et al*., 1983). The conventional method for sample preparation involves releasing intact nuclei by chopping fresh plant material in a compatible buffer (Doležel *et al*., 2007; Galbraith *et al*., 1983). Leaf material is always co-chopped with another plant species of known GS and ploidy, to calculate the relative difference in fluorescence and hence GS (Doležel *et al*., 2007; Galbraith *et al*., 1983; Temsch *et al*., 2022). GS is expressed and measured as a “C-value”, which is the entire DNA content of a nucleus (Greilhuber *et al*., 2005). The DNA content of a haploid nucleus, in its unreplicated state, is referred to as the ‘1C-value,’ and it is measured in units of picograms (pg) or million base pairs (Mbp) (Doležel *et al*., 2003; Greilhuber *et al*., 2005). One pg of DNA is equivalent to 978 Mbp (Doležel *et al*., 2003).

During fluorescence measurements of the samples with moderate levels of debris, it is recommended to capture data of 2000 events in total and 600 per sample peak to maintain relative standard error (SE) below 0.2 % (Koutecký *et al*., 2023). However, the conventional method of nuclei isolation is highly sensitive to plant chemistry, buffer chemistry, and chopping style (Loureiro *et al*., 2021; Loureiro *et al*., 2006a, b). Despite optimisation of the conventional method, some plant species remain recalcitrant to the production of the required number of nuclei (Koutecký *et al*., 2023; Loureiro *et al*., 2021; Temsch *et al*., 2022). In particular, plant material high in secondary metabolites can be challenging and often requires substantially different buffers and chopping styles to generate a repeatable result (Čertner *et al*., 2022; Loureiro *et al*., 2006a; Noirot *et al*., 2000). With a limited amount of sample, time, and other resources, recording this minimum number of events is a bottleneck for high throughput flow cytometry (Čertner *et al*., 2022).

It is becoming more common to use flow cytometry to estimate genome size prior to sequencing plant genomes (Doležel *et al*., 2007; Nakandala *et al*., 2023). In instances where plant materials are sourced from remote and geographically distant locations, the challenge arises in maintaining the freshness of the specimens over extended periods. The inherent difficulty in preserving plant material under such conditions renders it impractical for prolonged storage, consequently impeding the feasibility of GS estimation. The logistical constraints pose a significant obstacle to the preservation of plant material integrity, thereby limiting the scope and reliability of plant material for FCM (Čertner *et al*., 2022). A few studies have previously used fixed material, either ethanol preserved, paraffin fixed, or frozen material for ploidy and genome size estimation of plants (Bagwell *et al*., 1991; Cires *et al*., 2009; Dart *et al*., 2004; Halverson *et al*., 2008; Hopping, 1993; Jarret *et al*., 1995; Kolář *et al*., 2012; Nsabimana and Van Staden, 2006; Xavier *et al*., 2017). Despite this, frozen plant material is not generally used for flow cytometry in the estimation of ploidy and genome size (Čertner *et al*., 2022; Doležel *et al*., 2007). Little information is available about the relative fluorescence of the frozen and fresh.

Here we report a novel method using frozen plant material to release a high number of intact nuclei for accurate and precise estimation of the GS. This method is based on the homogenisation of frozen plant material by physical disruption (grinding and blending) and chemical disintegration of the cell wall (by detergents and buffers) to isolate intact nuclei (Sikorskaite *et al*., 2013; Workman *et al*., 2018; Zhang *et al*., 1995). The proposed nuclei isolation method can complement genome studies where plant material is frozen for long-term use. Furthermore, isolation of intact nuclei from frozen plant tissue can also be used to obtain high-quality genomic DNA for sequencing (Givens *et al*., 2011; Workman *et al*., 2018; Zhang *et al*., 1995).

Frozen plant material was previously reported to produce high debris content and low histogram resolution in attempts to estimate GS and ploidy (Čertner *et al*., 2022; Doležel *et al*., 2007). Moreover, a low yield of frozen nuclei has been considered a constraint for reliable estimation of the mean fluorescence (Hopping, 1993; Nsabimana and Van Staden, 2006). When assessing peak or histogram quality, the number of events within each peak and the coefficient of variance (CV) is considered (Doležel *et al*., 2007; Loureiro *et al*., 2021). A high level of debris compared to the fluorescence signal (i.e., low signal-to-noise ratio) can obscure the accuracy of the mean fluorescence peak and increase the CV (Doležel *et al*., 2007; Smith *et al*., 2018). The debris may contain DNA generated by rupturing cells and nuclei and, when incompatible with the buffer, can aggregate on the nuclei membrane to alter fluorescence readings (Čertner *et al*., 2022; Greilhuber *et al*., 2007; Nath *et al*., 2014). Secondary metabolites can also degrade the nuclear membrane and, therefore, interfere with the fluorescence readings (Doležel *et al*., 2007; Loureiro *et al*., 2021; Noirot *et al*., 2000). The presence of debris is common for many species (Čertner *et al*., 2022; Doležel and Bartoš, 2005). With nuclei isolation protocol from frozen plant material, the combination of several washing steps, use of buffers, filtration and centrifugations help to eliminate the debris in the form of intact cells and tissue residues (Workman *et al*., 2018).

The method reported here was applied to four plant species and fluorescence parameters were compared for the nuclei isolated from frozen and fresh preparations. The fluorescence data from both preparations was subjected to the conventional histogram analysis and debris compensated peak modelling approach to assess the accuracy of GS estimates.

## Materials and Methods

### Plant material

*Adenanthos sericeus* var. *sericeus* Labill.*, Hollandaea sayeriana* (F.Muell.) L.S.Sm., *Macadamia tetraphylla* L.A.S.Johnson and *Macadamia jansenii* C.L.Gross & P.H.Weston, representing approximately 2.5 times diversity in the GS, were selected for statistical comparison. These species were considered difficult due to the presence of polyphenols and tannins in the leaf material (Čertner *et al*., 2022; Gadea *et al*., 2022). Information about the chromosome number, ploidy and genome size estimates are available for *A. sericeus*, *H. sayeriana* (Jordan *et al*., 2015; Ramsay, 1963; Rao, 1957). Moreover, chromosome-level genome assemblies are available for *M. tetraphylla and M. jansenii* for determination of the accuracy of the genome size estimates (NCBI, 2023; Sharma *et al*., 2021). Young plants of *H. sayeriana* and *A. sericeus* were sourced from local nurseries and kept in glasshouse conditions at the University of Queensland. Leaf material for *M. tetraphylla* and *M. jansenii* was collected from Mt Coot tha Botanical Gardens, Brisbane. *Oryza sativa* subsp. *japonica* cv*. ‘*Nipponbare*’* (1C=388.8 Mbp/0.397 pg) was used as the internal standard and grown in glasshouse conditions at the University of Queensland (Project, 2005; Sasaki, 2005). *O. sativa* fulfilled the criteria of internal standards such as verified genome size stability, absence of anatomical or chemical features, and endopolyploidy impeding the fluorescence measurements (Project, 2005; Temsch *et al*., 2022). Additionally, *O. sativa* is easy to grow and maintain in glasshouse conditions in the study area.

### Pretreatment of leaf material

For frozen preparations, young, fully expanded, healthy leaves of the plants were collected in labelled perforated plastic bags and snap-frozen in liquid Nitrogen for 30 s. Subsequently, leaves were stored promptly at −80 °C until processed for nuclei isolation. For fresh preparations, fresh, fully expanded young leaves were sampled in a plastic bag with a moist paper towel and processed for nuclei isolation on the same day of the collection.

### Extraction of nuclei from fresh plant material

#### Buffers and reagents

- Modified Woody Plant Buffer (WPB) (as per Jordan *et al*., 2015; Loureiro *et al*., 2007a): 0.2 M Trizma hydrochloride (Sigma, 93363-50G), 0.04 M Magnesium chloride hexahydrate (Sigma, M2670-100G), 0.02 M EDTA.Na2 (Sigma, EA023-500G), 86 mM Sodium chloride (Sigma, 71380-500G), 10 mM Sodium metabisulfite (Sigma, S9000-500G), 1 % Triton X-100 (Chem Supply, TL125-P), UltraPure DNase/RNase free distilled water (Invitrogen, Cat. No. 10977-015, 300ml), 3% Polyvinylpyrrolidone −10 (Sigma, PVP10)
- Staining buffer (20 µl per 400 µl of sample): 100 µl Propidium Iodide (PI, 1mg/ml, Sigma, Product ID P4864-10ML), 1 µl of RNase 1 mg/ml. Keep the buffer on ice and cover it with aluminium foil due to light-sensitive nature of PI.

#### Equipment

47 mm diameter Petri dish (Advantec, Product ID PD-47A), single edge razor blades (Personna, Product ID 94-120-2), 40 µm polypropylene framed cell strainers (Biologix, Product ID 15-1040), 5 ml (12 x 75 mm) polystyrene round bottom tubes (Falcon, Product ID 0587866)

#### One step protocol to release nuclei from fresh plant material

For fresh preparations, one-step protocol was used as described in (Doležel *et al*., 2007). For each replicate, 40 mg of young fully expanded leaves of the test species co-chopped with 15 mg of the internal standard (*O. sativa)* in a petri dish using a single-edge razor blade in 500 µl ice-cold modified woody plant buffer (WPB) (Jordan *et al*., 2015; Loureiro *et al*., 2007a). The homogenate was filtered through a pre-soaked 40 µm nylon filter (Doležel *et al*., 2007). 20 µl of staining buffer (containing 100 µl of propidium Iodide (1 mg/ml) and 1 µl of RNase 1 mg/µl) was added to 400 µl of nuclei filtrate, and the sample was kept on ice until processed. Five biological replicates were performed for each species.

#### Protocol for Nuclei extraction from frozen leaf material

The nuclei isolation method of Workman *et al*. (2018) was opted as provided in Nuclei Isolation – LN2 Plant Tissue Protocol Document ID: NUC-LNP-001, Circulomics).

#### Reagents

Liquid N_2,_ spermidine trihydrochloride (Sigma, catalogue number S2501), spermine tetrahydrochloride (Sigma, cat. no. S1141), sucrose (Sigma, cat. no. S9378), Triton X-100 (Chem Supply, cat. No. TL125-P), polyvinylpyrrolidone-360 (Sigma, cat. no. PVP360), Trizma Base (Sigma, cat. no. T1503), potassium chloride (Scharlau, cat. no. 0401, 2-mercaptoethanol, 14 M (Sigma, cat. no. M3148), 0.5 M ethylene diamine tetra acetic acid (Biobasic Inc., Product: EB0185)

#### Equipments

Sterilised mortar and pestle, refrigerated centrifuge equipped with fixed angle rotor (Sigma Model 4-16K), Steriflip vacuum-driven filtration system with 20 µm nylon net filter (Merck Millipore, Cat. no. SCNY00020), 40 µm polypropylene framed cell strainers (Biologix, Product ID 15-1040), Pasteur pipettes (20 µl, 200 µl, 1000 µl, 5000 µl), 50 ml conical bottom centrifuge tubes (Corning, Product ID 430304), 1.5 ml microfuge tubes, 5 ml (12 x 75 mm) polystyrene round bottom tubes (Falcon, Product ID PID0587866)

#### Buffer preparation

- 10x Homogenisation buffer (HB): Trizma Base (0.1 M), Potassium chloride (0.8 M), ethylene diamine tetra acetic acid (0.1 M), spermidine (17 mM), spermine (17 mM), 10 M NaOH to adjust pH to 9. The solution can be stored in a glass bottle at 4 °C for up to one year.
- 100 ml Triton sucrose buffer (TSB): Triton X-100 (20 %), 10x HB (10 %), sucrose (0.5 M), Volume was made up to 100 ml with distilled water. The solution can be stored in a glass bottle at 4 °C for up to one year.
- 1000 ml 1x Homogenisation buffer (HB): 10x HB (10%), Sucrose (0.5 M), Volume was made up to 1 L with distilled water
- 50 ml/sample Nuclei Isolation Buffer (NIB): 1x HB (48.75 ml), TSB (1.25 ml), polyvinylpyrrolidone-360 (0.5 gm), Add 125 µl of 2-mercaptoethanol before use and keep NIB on ice.

#### Nuclear Isolation Protocol

1. Before starting the procedure of nuclei extraction, 50 ml of NIB per 2 gm sample was prepared fresh and stored at 4 °C.
2. 1.8 gm of the frozen leaf tissue of sample species and 0.2 gm of frozen leaf tissue of internal standard were taken in a sterilised, precooled mortar with liquid N_2_. Leaf material was submerged in liquid N_2._
3. The plant material was pulverised^1^ in a sterilised mortar and pestle in liquid N_2_. Hard leaves took longer to grind; therefore, jabbing converted big leaf parts into smaller pieces. After removing large chunks, small pieces were crushed into powder form with circular round motions of the pestle (Fig. S1A). This is a temperature-sensitive step; therefore, keep adding liquid Nitrogen to avoid thawing.
4. Homogenisation and nuclear isolation: Using a precooled spatula, leaf powder was quickly transferred to a precooled 50 ml falcon tube prefilled with 7.5 ml of NIB. Falcon tubes with 7.5 ml NIB were kept on ice before starting the procedure.
5. With 4-5 swirls, the powder was submerged in the NIB that no clumps were visible. Another 7.5 ml of NIB was added to the solution. The solution was gently mixed with the occasional end-to-end mixing for 2-3 min for 20 min. The solution was kept on ice to reduce the enzymatic activity of nucleases. After 20 min, a homogenate consisting of thousands of intact nuclei was ready (Fig. S1B).
6. Homogenate was filtered through 20 µm vacuum filtration system, and the filtrate was transferred in an empty 50ml falcon tube and kept on the ice (Fig. S1C, Fig. S1D).
7. Tubes were centrifuged at 7000 g and 4 °C for 20 min as the genome size was below 1000 Mbp. For large genomes (>1000 Mbp), centrifuge the tubes at 3000 g at 4 °C for 20 min. After centrifugation nuclei pellet was visible on the side or bottom of the tube; carefully discard the supernatant. At this stage, the pellet was green, representing contamination in the form of plant cell debris or secondary metabolites (Fig. S1E).
8. First wash: 7.5 ml of the NIB was added to the tube, and the pellet was mixed gently with a 10 ml pipette by pipetting out 7-10 times. Another 7.5 ml of NIB was added to the tube. The solution was kept on ice for 10 min with occasional gentle mixing, as in step 5. Tubes were centrifuged as indicated in step 6. After discarding the supernatant pellet should have a light colour (Fig. S1F).
9. Second wash: Added 10 ml of NIB and mixed the pellet gently with a 10 ml pipette by pipetting out 7-10 times. The tubes were kept in ice for 10 min with occasional gentle mixing. Tubes were centrifuged, as mentioned in step 7. The supernatant was discarded carefully after the centrifugation (Fig. S1G).
10. Final wash and aliquots: In the final wash, 7.5 ml NIB was added to the tube, and the pellet was mixed with a pipette, as indicated in step 8 (Fig. S1H). After mixing, the homogenate was equally allocated to five 1.5 ml microfuge tubes. Eppendorf tubes were centrifuged at 7000 g for 10 min. The supernatant was discarded carefully. A white nuclear pellet was visible at the bottom of the tubes. Nuclei pellets were snap frozen in liquid N_2_ after discarding the supernatant (no need to dry the tubes). Tubes were stored at −80 °C freezer until processed for FCM measurement.

#### Staining of frozen nuclei

Nuclei from frozen leaf material were isolated in pelleted form using the above protocol. Tubes containing frozen nuclei were kept on ice and 500 µl of ice-cold modified WPB was added to the ice-cooled 1.5 ml microfuge tubes. After five min, the nuclear pellet was mixed in the buffer 7-10 times using a P1000 pipette. The homogenate was filtered through a pre-soaked (in WPB) 40 µm nylon cell filter. 400 µl of the filtrate was added to the 5 ml round bottom tube, and 20 µl of the staining buffer (containing 100 µl of Propidium Iodide (Sigma: 1 mg/ml) and 1 µl of RNase 1 mg/µl) was added to the solution. The solution was mixed by flicking the bottom of the tube with finger 4-5 times, and tubes were kept on ice until loaded to flow cytometer.

#### Flow cytometry

Nuclei labelled with propidium iodide were excited by a blue laser (488 nm) and fluorescence was measured with a detector configured with a 695/40 nm bandpass filter on the Becton Dickinson LSR Fortessa X20 Cell Analyser. Fluorescence data was recorded on a linear scale of 256 channels (Koutecký *et al*., 2023). Leading trigger threshold was set to 5000. Fluorescence data was acquired for 20 min at a low rate (12 µl/min) which delivered 10-20 events/s for fresh preparations but 100-150 events/s for frozen preparations. Post-acquisition amplification of the signal was acquired by setting the forward scatter (FSC) detector voltage/gain to 320, side scatter (SSC) detector voltage to 179, and fluorescence detector voltage to 488 to position the internal standard peak at 1/5^th^ of the distance from the left end of the x-axis (Koutecký *et al*., 2023). Forward scatter and side scatter parameters were recorded on logarithmic scale and used to assist in

#### Conventional histogram analysis

For conventional histogram analysis, gating based on pulse analysis was used to separate single particles from aggregates in BD FACS DIVA software (v 8.0). Fluorescence pulse width on the y-axis was plotted against fluorescence pulse height on the x-axis to remove aggregates and debris (Fig. S2, S4, S6, S8, S10, S12, S14, S16). Although gating of the histogram is used to exclude the debris content from nuclei peaks assuming the Gaussian curve. However, considering the debris and the resolution of the histograms, our gating strategy involved selecting peaks in the middle of the population distribution, aiming to capture the most representative and homogenous portion of the population. The recommended limit of CV (i.e. < 5%) was also considered for gating of the histograms (Loureiro *et al*., 2007b). In addition, minimum requirements for accurate GS estimation were followed as 2000 events in total and 600 events per peak (Koutecký *et al*., 2023). However, despite the lower event count, the estimation of GS was still pursued for comparative purposes, acknowledging that the data obtained may provide valuable insights and contribute to the broader understanding of variations between the two methods of nuclei isolation and data analysis. GS was estimated (Eq. 1) in picograms (pg). 1pg was considered equivalent to 978 Mbp (Doležel *et al*., 2003). Apart from the GS, nuclei events per peak, debris %, and CV% were also recorded. The debris % was calculated (Eq. 2) to access the background debris (Nath *et al*., 2014).

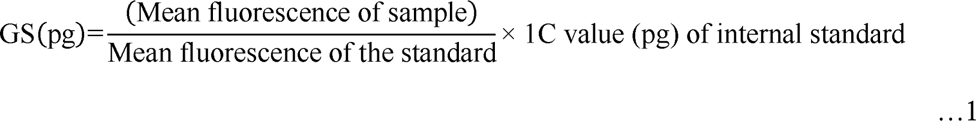

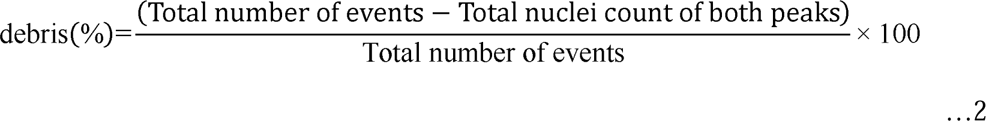

#### Histogram modelling and debris compensation based analysis

In this approach, data was subjected to peak modelling algorithms implemented in the ‘flowploidy’ package (v. 1.25.2) of R (v.4.2.3) (Ihaka and Gentleman, 1996; Smith *et al*., 2018). This modelling approach is based on the histogram-dependent non-linear least-squares algorithm for peak identification (Bagwell *et al*., 1991; Koutecký *et al*., 2023; Smith *et al*., 2018). Here, single nuclei events were isolated from aggregates and debris using the gating of the clusters of differential fluorescence based on particle size (Fig. S3, S5, S7, S9, S11, S13, S15, S17). Ratio of forward scatter pulse-height and fluorescence pulse height was plotted on the y-axis against fluorescence pulse-height on the x-axis to identify the single nuclei clusters (Fig. S3, S5, S7, S9, S11, S13, S15, S17). This package facilitated a non-linear regression function to fit a model, which was assessed for the goodness of fit based on residual Chi-Square (χ^2^) value (RCS) (Smith *et al*., 2018). After peak identification, data was processed with debris compensation accomplished through single-cut and multiple-cut algorithms implemented in the flowploidy package (Bagwell *et al*., 1991; Smith *et al*., 2018). RCS value between 0.7-4 for the best-fit model and recommended limits of CV (i.e. < 5%) were considered when gating to isolate debris and aggregates (Bagwell *et al*., 1991; Loureiro *et al*., 2007b; Sliwinska *et al*., 2022; Smith *et al*., 2018).

#### Experimental setup

For statistical comparison, data for four species (*A. sericeus*, *H. sayeriana, M. tetraphylla and M. jansenii*), two nuclei preparation methods (conventional method to extract nuclei from fresh material and proposed method to extract nuclei from frozen material), and two data analysis approach (debris compensated (including histogram modelling and debris compensation) and non-compensated (including conventional histogram analysis and no debris compensation)) were collected in one experiment.

#### Statistical analysis

Genome size was subjected to the three-way ANOVA (species x nuclei preparation x debris compensation) in ggplot2 (v 3.4.1) package of R. Post hoc comparisons were conducted using false discovery rate in R (v 4.2.3). Three-way interactions among species, method and compensation were tested for significance (CI-95%) on genome size and number of single nuclei events per peak. For the significance of the unequal variance, Levene’s test was used in R. Each combination of three variables was subjected to one-way ANOVA (CI-95%) coupled with post hoc comparison supported by false discovery rate correction. Analysis was conducted in the ‘ggstatsplot’ (v 0.11.0) and ‘ggplot2’ packages of R (v 4.2.3) (Patil, 2021; Wickham, 2011).

## Results

### Genome size estimates

There was a significant difference in genome size from the main effect of each factor, i.e., species, nuclei isolation method and debris compensation (Table 1, S1). Two-way interactions between ‘species and method’ (*p*=1.97e-06) and ‘species and compensation’ (*p*=0.00013) were significant (Table S1). However, three-way interaction among species, method, and compensation was not significant (*p*=0.51). One-way ANOVA was conducted for each combination of the three factors to find further significant interactions (Table S2).

**Table 1.**
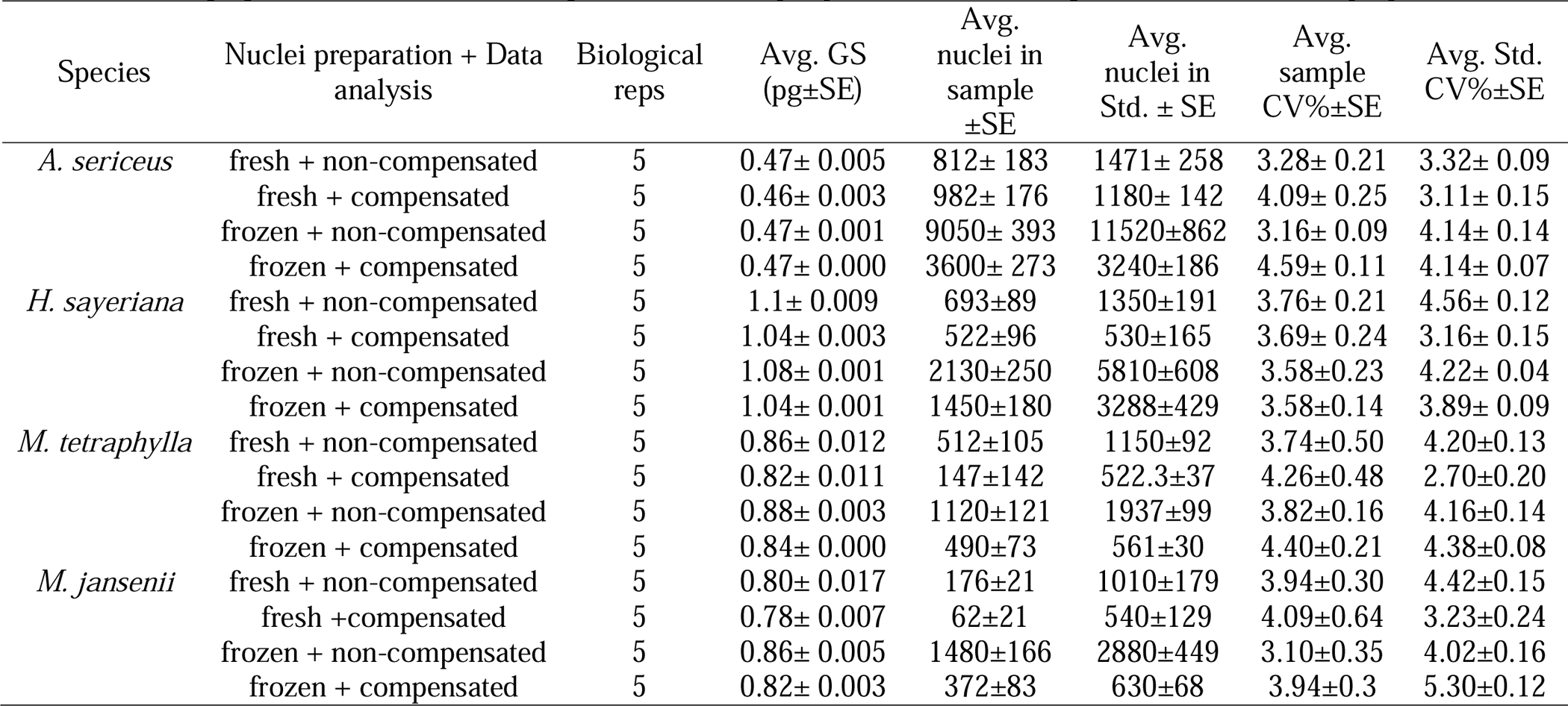
Average genome size estimates, single nuclei events per peak and CV% using fresh and frozen leaf preparations.

### Genome size estimates for *Adenanthos sericeus*

With conventional histogram analysis, no significant (*p*=0.21) difference was observed between the average 1C estimate of 0.47±0.001 pg from frozen nuclei and 0.47±0.005 pg from fresh nuclei (Fig. 1, Table 2S).

**Figure 1:**
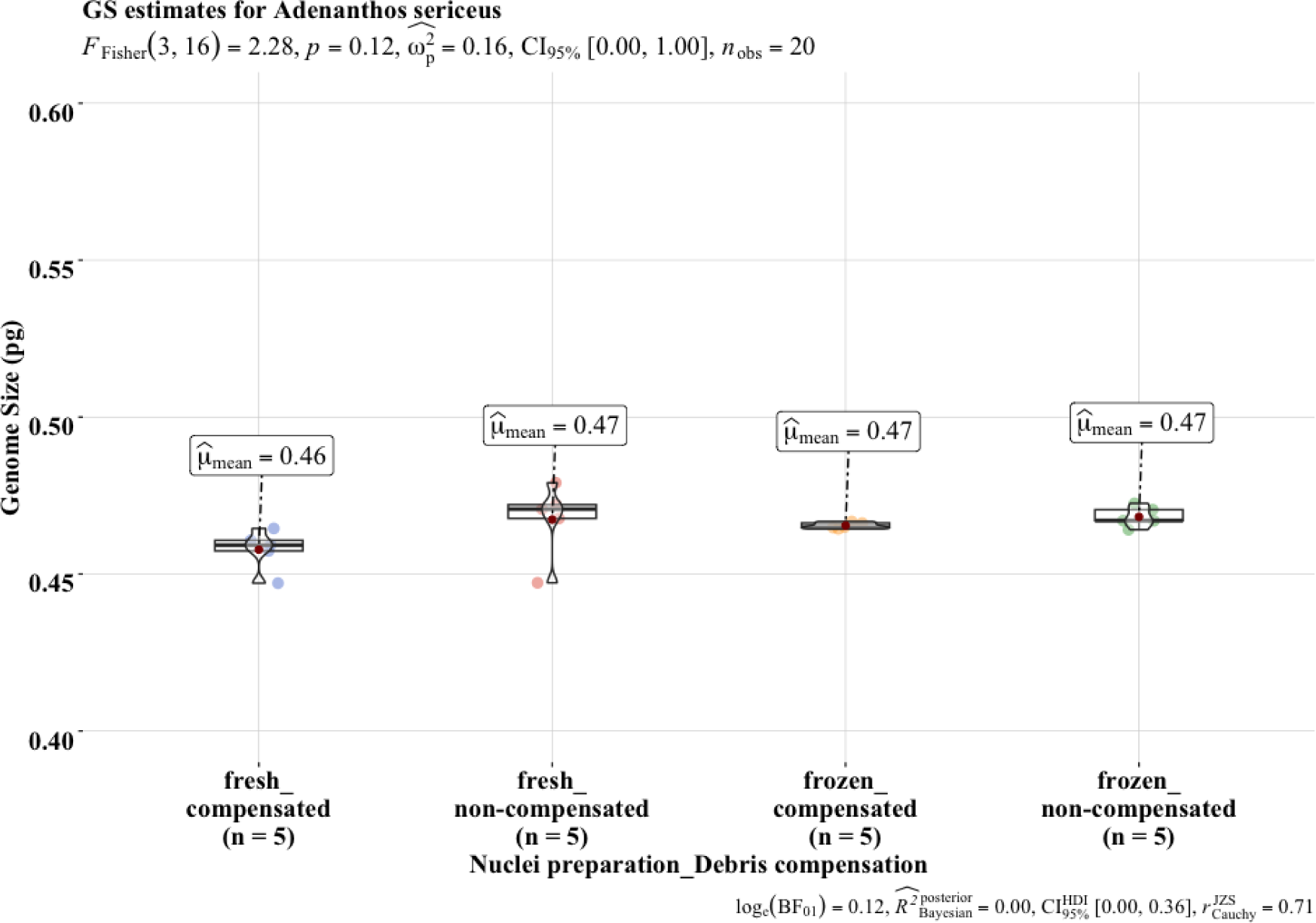
Comparison of the average 1C estimates for *A. sericeus* with different nuclei isolation and analysis approaches. Statistical analyses were conducted using Fisher’s ANOVA followed by a post hoc comparison using the false discovery rate (FDR) correction.

With model fitting and debris compensation, the average 1C estimate of 0.46±0.003 pg from fresh preparations was not significantly (*p*=0.14) different from the average estimate from conventional histogram analysis (Fig. 1). Similarly, the average 1C estimate of 0.47±0.000 pg from frozen nuclei and debris compensation was not significantly (*p*=0.21) different from the average estimate from conventional histogram analysis (Fig. 1).

### Genome size estimates for *Hollandaea sayeriana*

With conventional histogram analysis, the average 1C estimate of 1.09±0.001 pg from frozen nuclei was not significantly (*p*=0.21) different from 1.10±0.009 pg of fresh preparations (Fig. 2).

**Figure 2:**
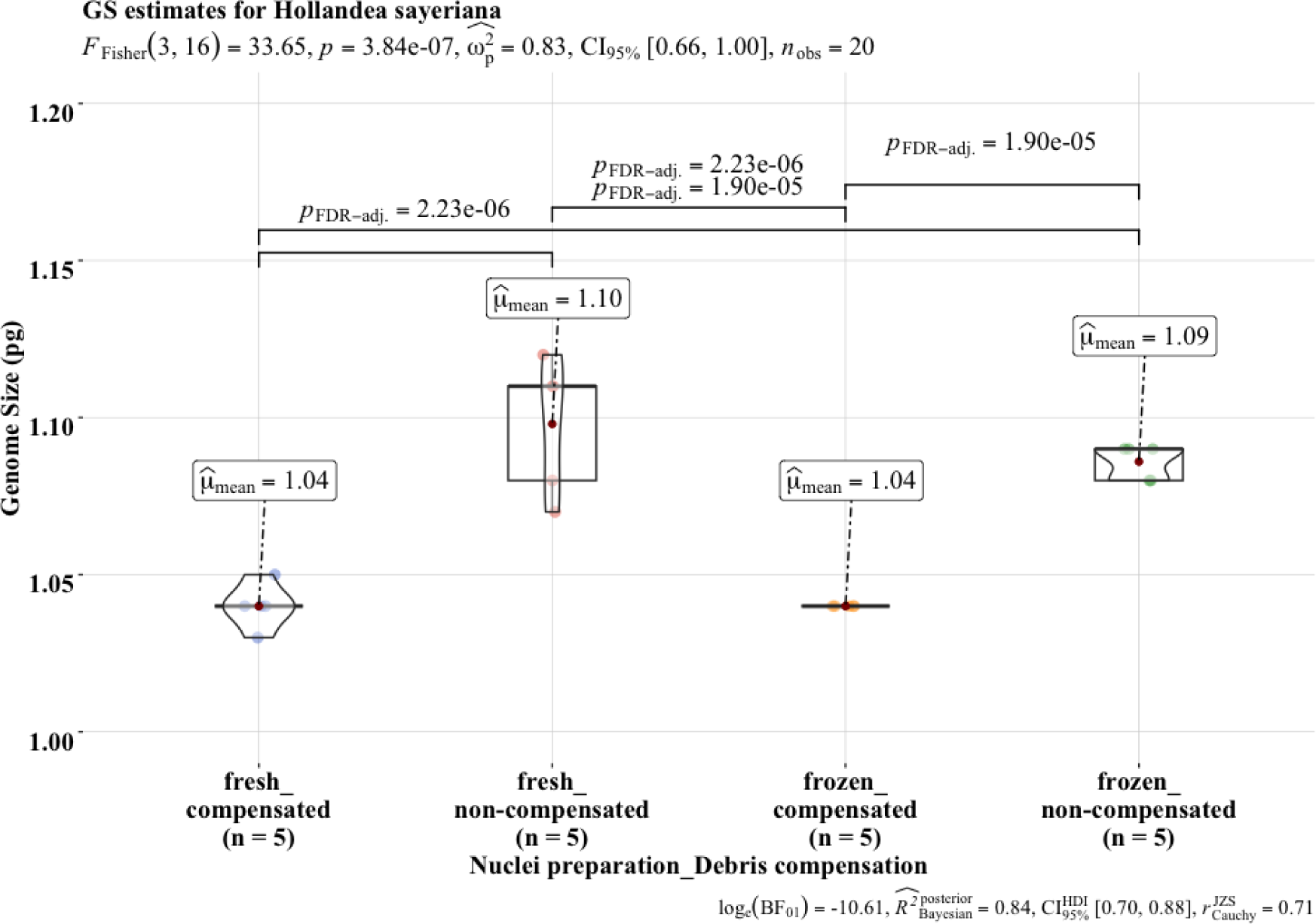
Comparison of the average 1C estimates for *H. sayeriana* using the difference nuclei isolation and analysis approaches. Statistical analyses were conducted using Fisher’s ANOVA followed by a post hoc comparison using the false discovery rate (FDR) correction.

With peak modelling and debris compensation, the average GS estimate of 1.04±0.001 pg from frozen nuclei was significantly lower (*p*=1.90e-05) than the average estimate from conventional histogram analysis of 1.08±0.001 pg (Fig. 2). Similarly, the average 1C estimate of 1.04±0.003 pg from fresh nuclei was significantly smaller (*p*=2.23e-06) than the average 1C estimate of 1.10±.009 pg without peak modelling and debris compensation. The GS estimate from fresh and frozen preparations was not significantly (*p*=1) different (Fig. 2). Although the average 1C value of 1.04±0.001 pg from frozen preparations and debris compensation was more precise (*p*=0.1, CI=90%) than estimates from other methods.

### Genome size estimates for *Macadamia tetraphylla*

With conventional histogram analysis and without debris compensation, the average 1C estimate of 0.86±0.012 pg fresh preparation was not significantly (*p*=0.22) different from the average 1C estimate of 0.88±0.003 pg from frozen preparations (Table 2S, Fig. 3).

**Figure 3:**
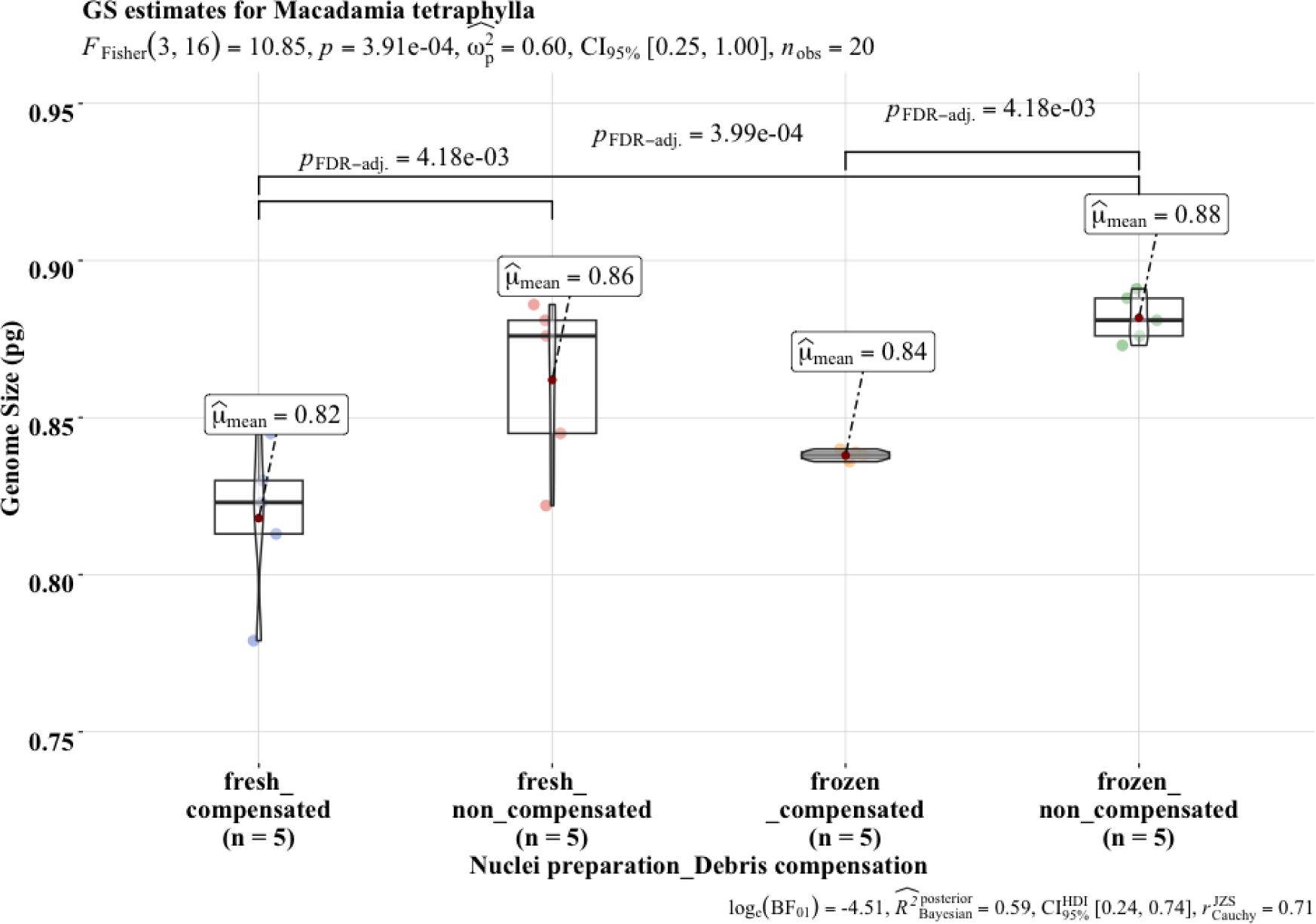
Comparison of average 1C estimates for *M. tetraphylla* derived using different nuclei isolation and analysis approaches. Statistical analyses were conducted using Fisher’s ANOVA followed by a post hoc comparison using the false discovery rate (FDR) correction.

With peak modelling and debris compensation, the average 1C estimate of 0.82±0.011 pg from fresh preparation was not significantly different (*p*=0.058) from the average 1C estimate of 0.86±0.012 pg with non-compensated data (Fig. 3). With debris compensation, the average 1C estimate of 0.84±0.000 pg from frozen preparations was significantly lower (*p*=4.18e^−03^) than the average 1C estimate of 0.88±0.003 pg from conventional histogram analysis (Fig. 3). The GS estimate from frozen nuclei and debris compensation was not significantly (*p*=0.15) different to the 1C estimate from fresh nuclei and debris compensation (Fig. 3).

### GS estimates for Macadamia jansenii

With conventional histogram analysis, the average 1C estimate of 0.86±0.005 pg from frozen preparations was significantly higher (*p*=1.40e-03) than the average 1C estimate of 0.80±0.017 pg from fresh preparations (Fig. 4, Table 2S).

**Figure 4:**
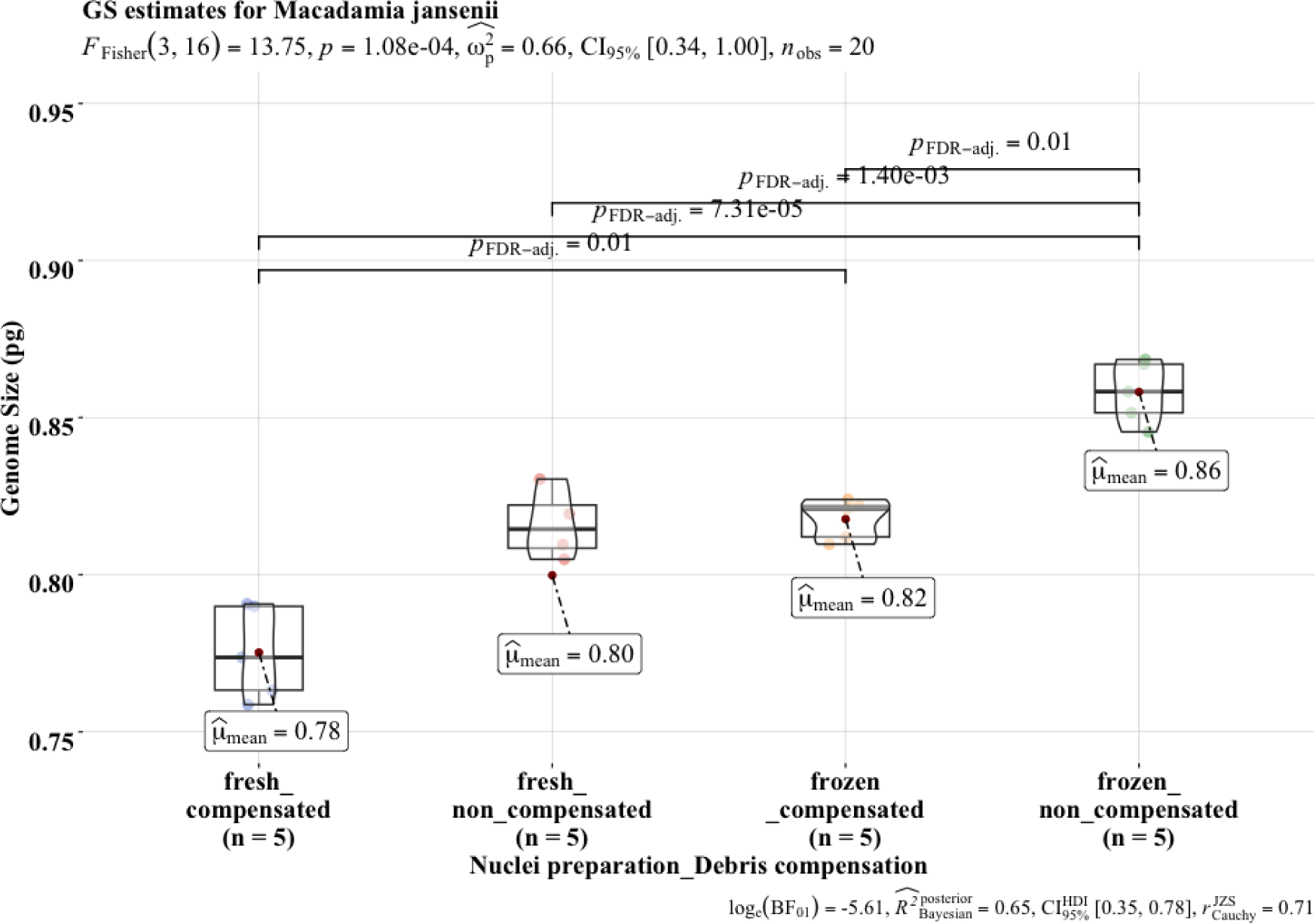
Comparison of the average 1C estimates for *M. jansenii* derived using different nuclei isolation and analysis approaches. Statistical analyses were conducted using Fisher’s ANOVA followed by a post hoc comparison using the false discovery rate (FDR) correction.

With peak modelling and debris compensation, the average 1C estimate of 0.82±0.003 pg from the frozen preparations was significantly (*p*=0.01) lower than the average 1C estimate of 0.86±0.005 pg from non-compensated analysis (Fig. 4). Whereas the average 1C estimate of 0.78±007 pg from fresh preparations and debris compensation was not significantly (*p*=0.32) different from the average estimate of 0.80±0.017 pg from fresh preparation and conventional histogram analysis (Fig. 4, Table 2S).

### Single nuclei event count per peak

The main effects of species, method and debris compensation were significant (*p*<0.001) for single nuclei event in the peak of test species (Table S3) and internal standard (Table S5). All two-way interactions for single nuclei count per peak were significant (*p*<0.001) (Table S3-, S5). Three-way interaction among species, method and debris compensation were significant for internal standard (*p*=2.77e-11) and test species (*p*=1.86e-16). One-way ANOVA was conducted for each combination of the three factors to find further significant interactions for nuclei count per peak (Table S4, S6).

### Single nuclei event count per peak for *Adenanthos sericeus*

With conventional histogram analysis, the average single nuclei count for *A. sericeus* peak with frozen preparations was 9050±393 (Table 1). This was significantly (*p*=1.29e-12), approximately nine times higher than the average of 982±176 events from fresh preparations (Fig. 5, Table 1). Similarly, for the *O. sativa* peak, the average single nuclei count was 11520±862 with frozen preparations. This was significantly (*p*=1.76e-10), approximately eight times, higher than the average of the fresh preparations (Fig. 5).

**Figure 5:**
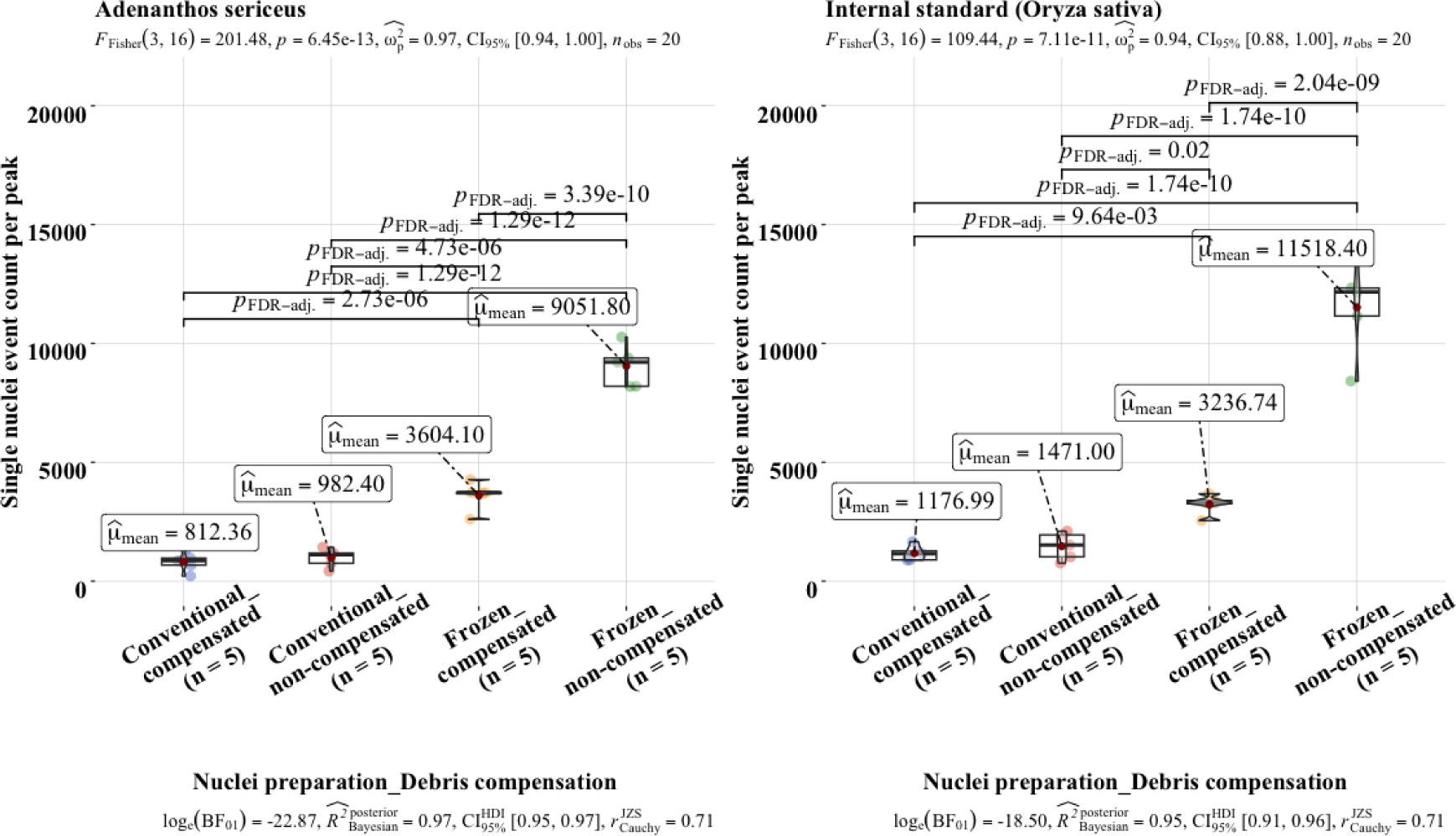
Comparison of average single nuclei event count per peak for experiments on *A. sericeus* and *Oryza sativa* ssp japonica var.’Nipponbare’. Statistical analyses were conducted using Fisher’s ANOVA followed by a post hoc comparison using the false discovery rate (FDR) correction.

With peak modelling and debris compensation, the average single nuclei count for *A. sericeus* reduced significantly (*p*=3.39e-10) to 39.8 % of the original events (Fig. 5, Table 1). Likewise, the single nuclei count for the internal standard decreased significantly (*p*=2.04e-09) to 28.1 % of the original events after model fitting and debris compensation (Fig. 5).

Despite significant reduction after debris compensation, the average events of 3600±273 for *A. sericeus* with frozen preparations were significantly (*p*=2.73e-06) higher than the average of 812±183 from fresh preparation and debris compensation (Fig. 5). The average nuclei events of 3240±186 for the peak of *O. sativa* from the frozen preparations was significantly (*p*=9.64e-03), approximately three times higher than the average of 1180±142 events from fresh preparations (Fig. 5).

### Single nuclei event count per peak for *Hollandaea sayeriana*

For non-compensated data from frozen nuclei preparations, the average single nuclei event count for *H. sayeriana* peak was 2130±250 (Table 1). This was significantly (*p*=4.75e-05), and nearly three times higher than the average of 693±89 from fresh preparations (Fig. 6, Table 1). The *O. sativa* peak had an average of 5810±608, which was significantly (*p*=1.59e-06), above four times higher than the average of 1350±191 events from fresh preparations (Table 1, Fig. 6).

**Figure 6:**
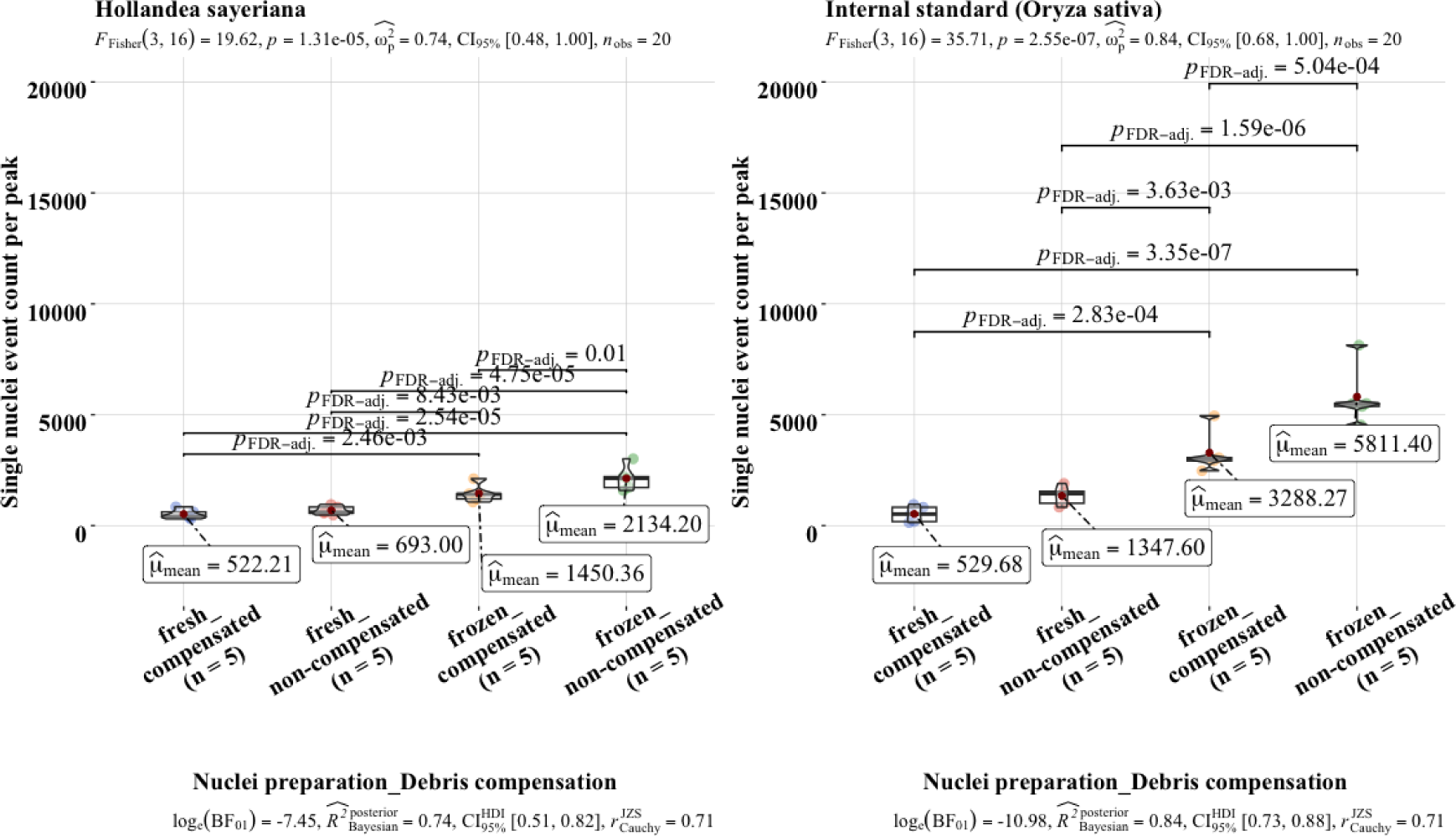
Comparison of average single nuclei event count per peak for experiments on *H. sayeriana* and *Oryza sativa* ssp. japonica var.’Nipponbare’. Statistical analyses were conducted using Fisher’s ANOVA followed by a post hoc comparison using the false discovery rate (FDR) correction.

With peak modelling and debris compensation, the average single nuclei count of 1450±180 for *H. sayeriana* peak from frozen preparations was significantly (*p*=2.46e-03), nearly three times higher than the average of 522±96 from fresh preparations (Table1, Fig. 6). Single nuclei event count for the internal standard peak with the frozen preparations reduced significantly (*p*=5.04e-04) by 43.4 % after data compensation (Table 1, Fig. 6).

### Single nuclei event count per peak for *Macadamia tetraphylla*

With conventional histogram analysis, the average single nuclei count of 512±105 for *M. tetraphylla* peak from fresh preparations was significantly lower (*p*=1.48e-05), nearly half of the average of 1120±121 from fresh preparations (Fig. 7). The average count of 1937±99 for the *O. sativa* peak from frozen preparation was significantly (*p*=1.72e-06) higher than the average of 1150±92 from fresh preparations (Fig. 7, Table 1).

**Figure 7:**
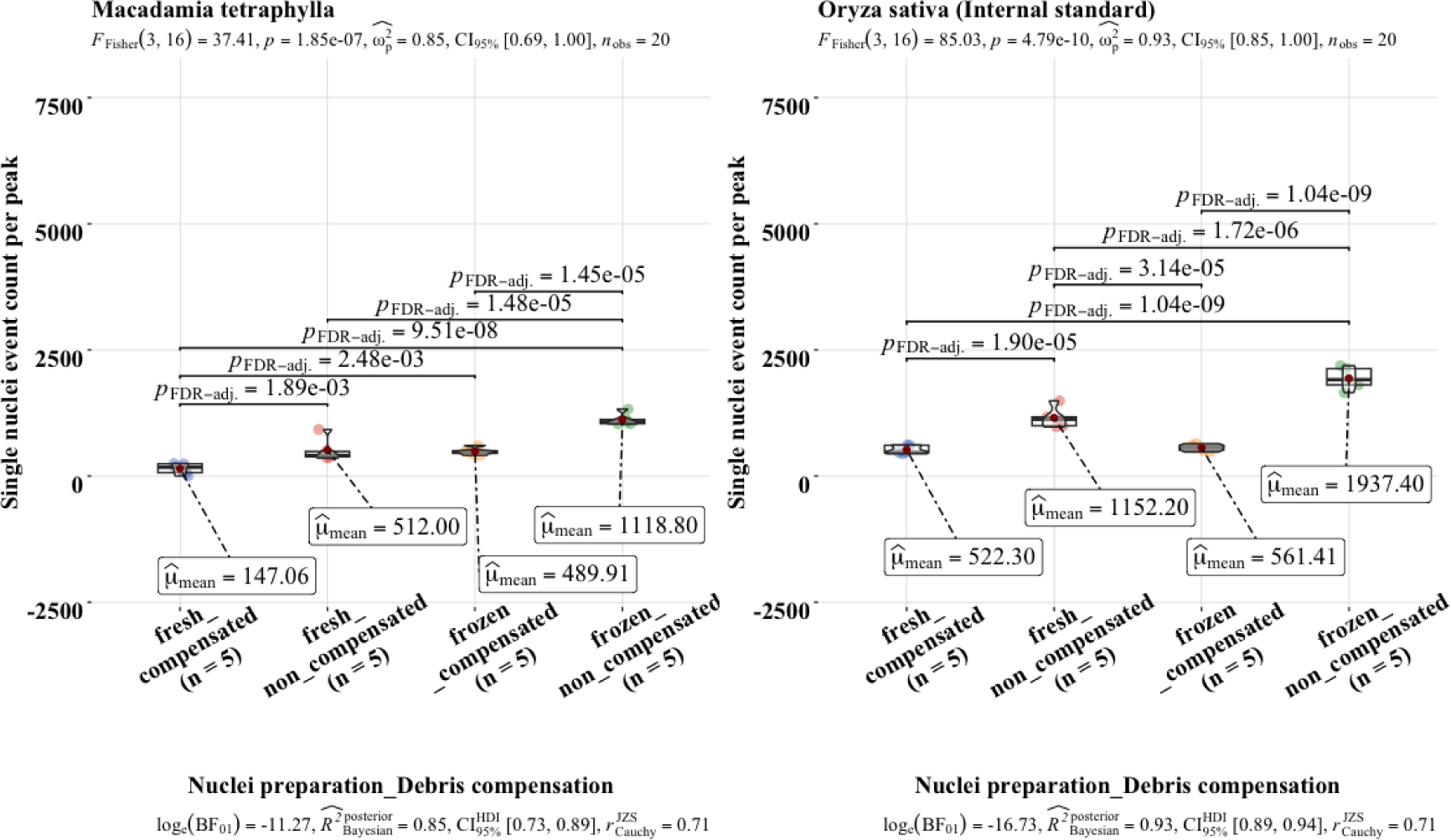
Comparison of the average single nuclei event count per peak experiments on *M. tetraphylla* and *Oryza sativa* ssp. japonica var.’Nipponbare’. Statistical analyses were conducted using Fisher’s ANOVA followed by a post hoc comparison using the false discovery rate (FDR) correction.

After peak modelling and debris compensation, average single nuclei for *M. tetraphylla* peak reduced significantly (*p*=1.45e-05) to 490±73. Despite the reduction, the average count from frozen preparations was significantly (*p*=2.48e-03), higher than the average of 147±14 from fresh preparations (Fig. 7).

### Single nuclei event count per peak for *Macadamia jansenii*

With conventional histogram analysis, the average single nuclei count of 1480±166 for *M. jansenii* frozen preparations was significantly (*p*=5.03e-08), over eight times higher than the average of 176±21 from fresh preparations (Fig. 8). Similarly, for the *O. sativa* peak, nuclei count with frozen preparation was 2880±449, which was significantly (*p*=6.01e-06), nearly three times, higher than the average of 1010±179 from fresh preparations (Fig. 8, Table 1).

**Figure 8:**
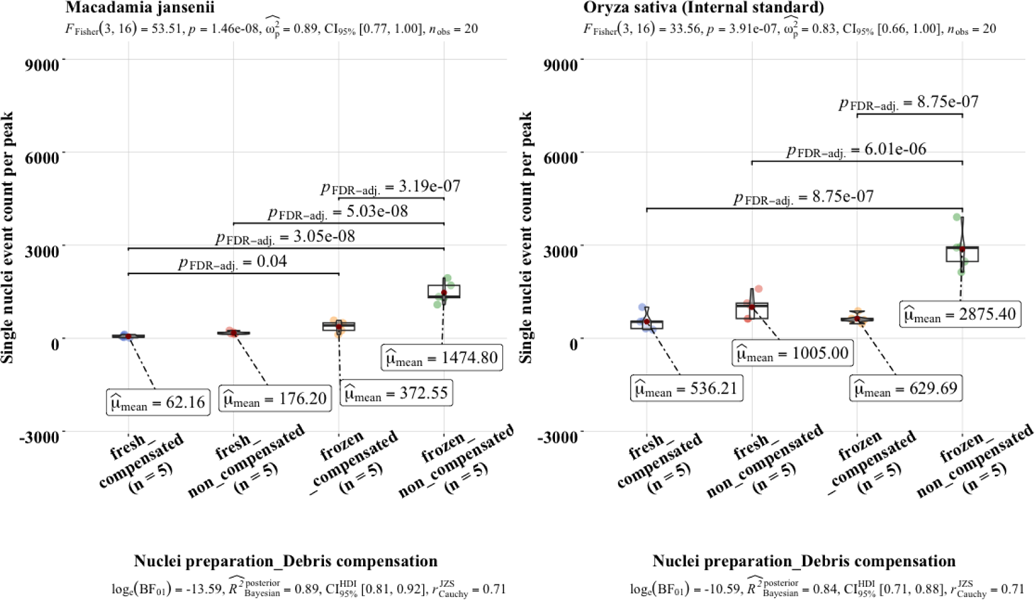
Comparison of single nuclei event count per peak for experiments on *M. jansenii* and *Oryza sativa* ssp. japonica var.’Nipponbare’. Statistical analyses were conducted using Fisher’s ANOVA followed by a post hoc comparison using the false discovery rate (FDR) correction.

After peak modelling and debris compensation, the average nuclei count for *M. jansenii* reduced significantly (*p*=3.19e-07) to 372±83 from frozen preparations. The average count for *O. sativa* peak also reduced significantly (*p*=8.75e-07) to 630±68 in frozen preparations (Fig. 8). Despite the significant reduction, the average count of *M. jansenii* peak in frozen preparations was significantly (*p*=0.04), higher than the average count from fresh preparation. The average count for *O. sativa* peak after debris compensation was not significantly (*p*=0.66) different in frozen and fresh preparations (Fig. 8).

### Debris and background noise

For samples representing a low signal-to-noise ratio, separation of the intact nuclei from the debris particles is difficult with conventional histogram analysis that inherently excludes debris compensation. The debris particles are also counted as a single nuclei event when a histogram is drawn around the fluorescence peak. Therefore, counting the exact number of intact nuclei with high background noise is not possible. The debris factor was estimated as the proportion of total single nuclei against all events. Except for *M. tetraphylla*, fresh preparations had significantly (*p*<0.001) higher background noise than frozen preparations (Fig. 9). For *M. tetraphylla,* debris was similar in fresh and frozen preparations (Fig. 9).

**Figure 9:**
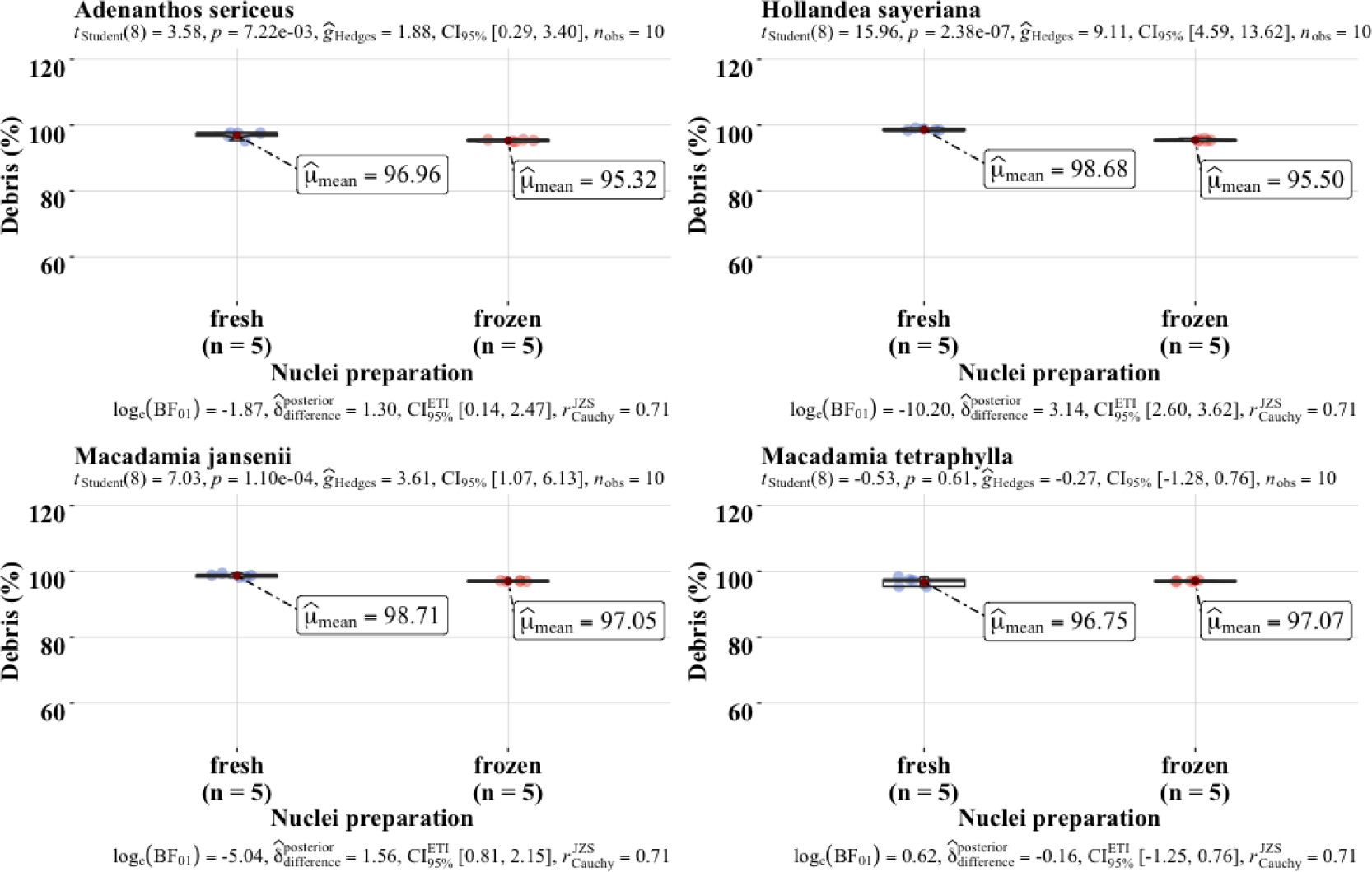
Comparison of the debris factor for all four species for fresh and frozen nuclei preparations. Statistical analyses were conducted using student t-test.

The average cumulative nuclei yield for both *A. sericeus* and *O. sativa* peaks were nearly 91% higher in frozen preparations, with 20570 nuclei events in both peaks (Table 1). Whereas the average debris content of 95.3±0.13 % for frozen preparations was significantly *(p*=7.22e-03) lower than the average of 97±0.44 % from fresh preparations (Fig. 9).

For *Hollandaea sayeriana*, the nuclei yield for *H. sayeriana* and *O. sativa* peak was 7500 in frozen preparations, nearly four times higher than that of fresh preparations. The average debris content of 95.5±0.14 % from frozen preparations was significantly (*p*=2.38e-07) lower than the average debris of 98.7±0.14 % from fresh preparations (Fig. 9).

For *Macadamia jansenii*, the average cumulative single nuclei yield for both *M. jansenii* and *O. sativa* peaks was 4350 in frozen preparations, nearly four times higher than in fresh preparations. The average debris content of 97.0±0.04 % from frozen preparations was significantly (*p*=1.10e-04) lower than the average of 98.7±0.23 % from fresh preparations (Fig. 9).

For *Macadamia tetraphylla*, the average event yield was 3060 in frozen preparations. However, the yield in fresh preparations was 1660, approximately two times lower than in frozen preparations. The average debris of 97.0±0.10 % was slightly (*p*=0.61) higher than the average debris factor of 96.8±0.59 % from fresh preparations (Fig. 9).

## Discussion

Rapid progress in genome sequencing, coupled with applications in plant breeding and cytological studies, has promoted the implementation of FCM as a complementary approach (Galbraith *et al*., 2021). However, due to the limitations of the plant material storage, preservation strategies and absence of immediate FCM analysis, GS estimates for several species cannot be estimated (Čertner *et al*., 2022; Greilhuber *et al*., 2007). In addition, the conventional nuclei isolation from fresh material remains ineffective in recalcitrant species for several reasons (Doležel *et al*., 2007; Loureiro *et al*., 2021; Temsch *et al*., 2022). Here, we presented an approach of using frozen plant material to release intact nuclei, which can be complemented with histogram modelling-based analysis to estimate the genome size showing no difference from the fresh preparations. GS estimates from frozen plant material can be a good asset for genome sequencing studies where the sample is frozen after retrieval, or fresh plant material is not available. After genome estimates, isolated nuclei can be used for the high molecular weight DNA extractions (Givens *et al*., 2011; Workman *et al*., 2018).

Nuclei isolated from frozen leaf material with conventional chopping method have been used previously for genome size and ploidy estimation in plants (Dart *et al*., 2004; Halverson *et al*., 2008; Nsabimana and Van Staden, 2006). However, nuclei isolation from fixed tissue has always been debatable and often rejected due to chromatin restructuring from fixative agents (Greilhuber *et al*., 2007; Xavier *et al*., 2017). Although a few studies have observed intact nuclei despite several steps of mechanical and chemical disintegration of the frozen tissue (Givens *et al*., 2011; Jiao *et al*., 2012; Sikorskaite *et al*., 2013; Zhang *et al*., 1995). Moreover, studies have also noticed the intactness of nuclei after the freezing/thawing process (Hopping, 1993; Kratochvílová *et al*., 2019). The similarity of GS estimates from frozen and fresh preparations in this study suggests the intactness of the frozen nuclei. The integrity of the frozen nuclei within a cell is affected by thawing of ice crystals in extracellular and intracellular fluid (Kratochvílová *et al*., 2019). In the proposed method, however, the grinding process is conducted in liquid Nitrogen to maintain the structural integrity. Blenders and pulverisers have been used for grinding frozen material (Sikorskaite *et al*., 2013). However, Zhang *et al*. (1995) suggested the use of mortar and pestle for a higher yield of intact nuclei. In addition, Givens *et al*. (2011) also opted a similar way of isolating intact nuclei from frozen fungal material. The thawing damage is largely dependent on the thawing conditions (Kratochvílová *et al*., 2019). Therefore, the nuclei isolation process is conducted at lower temperatures to reduce the enzymatic activities and thawing process.

In earlier studies, frozen plant samples produced a low signal-to-noise ratio, and results were disregarded (Cires *et al*., 2009; Doležel *et al*., 2007; Greilhuber *et al*., 2007; Xavier *et al*., 2017). Some studies also produced histograms of reasonably high resolution, representing the potential of frozen plant material for estimation of the preliminary or absolute estimates of ploidy and genome size (Čertner *et al*., 2022; Cires *et al*., 2009). Nonetheless, it was believed that a probable alteration in fluorescence might lead to a deviation in the genome size estimations. The similarity of the GS estimates from frozen nuclei with the estimates from fresh nuclei in this study indicates that the two methods can be used interchangeably to estimate the absolute DNA content (Table 2). In addition, nuclei isolation protocol from frozen plant material is optimised for reducing the impact of the secondary metabolites for high-quality DNA extractions (Cushman, 1995; Workman *et al*., 2018; Zhang *et al*., 1995).

**Table 2.** Genomesize (GS) estimates using fresh and frozen leaf preparations against reference genomes (Clark *et al*., 2016; NCBI, 2023; Sayers *et al*., 2022; Sharma *et al*., 2021).

| Species | Preparation + Debris compensation | Replicates | GS (pg) | GS (pg, Whole genome sequence) |
| --- | --- | --- | --- | --- |
| *Macadamia tetraphylla* | fresh + non-compensated | 5 | 0.86±0.0123 | 0.81 |
| *Macadamia tetraphylla* | fresh + compensated | 5 | 0.82±0.011 | 0.81 |
| *Macadamia tetraphylla* | frozen + non-compensated | 5 | 0.88±0.003 | 0.81 |
| *Macadamia tetraphylla* | frozen + compensated | 5 | 0.84±0.000 | 0.81 |
| *Macadamia jansanii* | fresh + non-compensated | 5 | 0.80±0.017 | 0.80 |
| *Macadamia jansanii* | fresh + compensated | 5 | 0.78±0.007 | 0.80 |
| *Macadamia jansanii* | frozen + non-compensated | 5 | 0.86±0.005 | 0.80 |
| *Macadamia jansanii* | frozen + compensated | 5 | 0.82±0.003 | 0.80 |

### Integration of model fitting and debris compensation algorithms complements frozen preparations for higher precision and accuracy

Plant species without significant debris can have high-resolution fluorescence peaks if the sample is prepared with the best practices (Loureiro *et al*., 2021). The high resolution of the peaks allows robust estimation of the genome size with the conventional histogram analysis (Koutecký *et al*., 2023). However, with recalcitrant plant species, debris amount can be challenging for a reliable GS estimation (Koutecký *et al*., 2023; Loureiro *et al*., 2006a). Data analysis, particularly debris compensation, of FCM data can affect the GS estimates significantly (Wersto and Stetler-Stevenson, 1995).

For *M. tetraphylla*, without peak modelling and debris compensation, the GS estimate from frozen nuclei was 0.88±0.003 pg. This was not significantly (*p*=0.22) different from the 0.86±0.0123 pg of fresh preparations. However, peak modelling and debris compensation reduced the average GS estimate from fresh and frozen preparations significantly. The average 1C estimate of 0.84±0.000 pg from frozen preparations was not significantly (*p*=0.15) different than that of 0.82±0.011 pg from fresh preparations. After peak modelling and debris compensation, the GS estimate matched closely to the reference genome size of 0.81 pg (Clark *et al*., 2016; NCBI, 2023; Sayers *et al*., 2022) (Table 2).

The average 1C estimate of 0.86±0.005 pg of *M. jansenii* with conventional histogram analysis of the frozen preparations was significantly (*p*=0.01) higher than the average estimate of 0.82±0.003 pg from peak modelling and debris compensation. The average GS estimate from frozen preparations and debris compensated analysis corresponded with the reference genome size of 0.80 pg (Clark *et al*., 2016; NCBI, 2023; Sharma *et al*., 2021) (Table 2). However, given the limitations of the sequencing and assembling pipelines and genome complexity, genome assemblies can have gaps (Gladman *et al*., 2023; Suda and Leitch, 2010). Moreover, the genome assemblies exclude the endosymbiotic organellar DNA. Likewise, a slight error in fluorescence can also bring erroneous estimate of the genome size with flow cytometry.

With conventional histogram analysis, the GS estimates were significantly higher for all except *A. sericeus.* Peak modelling and debris compensation slightly improved the precision for all species. For *M. tetraphylla and M. jansenii,* the average GS estimates from frozen preparations were more accurate after peak modelling and debris compensation. However, the accuracy of the estimates for *A. sericeus* and *H. sayeriana* could not be verified in the absence of a reference genome sequence.

Lower precision from the fresh preparations could be due to variations in leaf chemistry, sample processing and handling errors each time a replicate was prepared. Nuclei isolation protocol from frozen material is designed to remove cytosolic contaminants that can hinder the fluorochrome binding and number of intact nuclei. Therefore, the debris occurring in data from this method is assumed primarily to be nuclear debris generated from subsequent nuclei damage by single cut or multiple cuts (Bagwell *et al*., 1991; Smith *et al*., 2018). However, non-nuclear debris can be other metabolites of the plant cells that can affect the fluorescence (Loureiro *et al*., 2021).

### High nuclei yield with frozen methodology increases the statistical significance of GS estimates

Previous attempts to use frozen leaf material for genome size and ploidy estimation employed the conventional chopping approach in a suitable buffer to release nuclei (Cires *et al*., 2009; Dart *et al*., 2004; Halverson *et al*., 2008; Nsabimana and Van Staden, 2006). However, low nuclei counts were observed when frozen plant material was chopped using the conventional method for ploidy determination (Nsabimana and Van Staden, 2006). Meanwhile, the proposed method of homogenisation of frozen plant material, with significantly higher nuclei yield, can outweigh the conventional approach of nuclei isolation for some challenging species. A large number of nuclei per peak reduce the data variability, subsequently improving accuracy by limiting the effects of outliers. With a small number of events, high variability in fluorescence can generate erroneous average fluorescence. In fresh preparations, slight variations in the tissue type, chopping and handling can bring fluorescence and GS variances when preparing a biological replicate. Such variations associated with leaf chemistry and handling can be avoided in the frozen preparations as multiple replicates can be prepared in one process. Frozen leaf material can be stored at controlled temperature for flow cytometry and genome sequencing purposes.

The low nuclei yield from fresh preparations of *Macadamia tetraphylla* and *M. jansenii* was a significant drawback of the fresh preparations of nuclei. This could be due to the incompatibility of the buffer or the recalcitrance of the species. For both *M. tetraphylla* and *M. jansenii*, the nuclei counts per peak were less than the minimum requirements of 600 events per peak for accurate GS estimation (Koutecký *et al*., 2023). Estimates from a small number of events represent a high vulnerability to outliers and consequently represent more variability. Moreover, due to the low signal-to-noise ratio, peaks with fewer events are obscured by debris. In large-scale studies including several challenging species, nuclear isolation from frozen material can be far more productive than fresh preparations. Although the amount of tissue used in frozen preparations is almost 10 times higher than in fresh preparations. However, chopping and homogenising a large quantity of leaf material is challenging, and it can negatively affect the quality of isolated nuclei by adding more contaminants and debris. The impacts of contaminants are well described (Loureiro *et al*., 2006b). In frozen preparations, disruption and homogenisation of the large quantity of leaf material can be achieved by maintaining the intactness of the nuclei under cryogenic conditions. In addition, several washes with buffer facilitate the removal of contaminants and make it processible for model fitting, debris compensation, and other statistical analyses.

For genome sequencing applications, inaccurate GS estimates may enhance the overutilisation of the resources, and this can reduce the applicability of the genome sequence applications. In the absence of fresh plant material or an effective nuclei isolation method, genome size estimation for several plant species is challenging. Often, an estimate from the nearest species is used as a guide for approximation. However, genome estimates from the different plants, sibling species, subspecies or cryptic species can represent significantly different genome sizes. Ideally, the estimation of GS of the plant to be sequenced should be from the same plant or plants. The methodology proposed here can be used where a genome size estimate is required for the plant to be sequenced and plant material is available in frozen form. The frozen material is widely used to release the intact nuclei for DNA extraction. The fluorescence of the frozen nuclei can be complemented with histogram modelling for precise and accurate estimation of the genome size. GS estimates from the frozen nuclei are similar to estimates from the fresh nuclei.

The plant material in this study was snap-frozen and processed within a short time span. However, the literature claims decay in sample quality over time that can potentially cause lower histogram resolution and shift of fluorescence. Storage strategies can be explored further. In addition, further studies can be conducted to investigate the effect of different compositions of the buffers and cryoprotectants on frozen preparations. In trial experiments, a difference in the debris and number of single nuclei events was observed when different buffers were compared for the scatter of the peaks (data not presented here).

## Supplementary Data

The following supplementary data are available at JXB online.

Supplementary Tables S1-S6

*Table S1*. Results of three-way interaction for species, nuclei extraction method and debris compensation in genome size estimation.

*Table S2*. Results of one-way ANOVA for all combinations of species, nuclei extraction method and debris compensation on genome size estimation.

*Table S3.* Three-way interactions among species, nuclei isolation method, and debris compensation on the single nuclei event count for the test species.

*Table S4.* Results of one-way ANOVA for all combinations of species, nuclei extraction method and debris compensation on nuclei events in the peak of sample species.

*Table S5.* Three-way interactions among species, nuclei isolation method, and debris compensation on single nuclei event count for internal standard species

*Table S6.* Results of one-way ANOVA for all combinations of species, nuclei extraction method and debris compensation on nuclei events in the peak of standard species.

Supplementary Figures S1-S17

*Fig. S1.* The process of nuclei isolation from frozen leaf material.

*Fig. S2.* Conventional histogram analysis for the fluorescence data of fresh preparations of *Oryza sativa* (P1) and *Adenanthos sericeus* (P2).

*Fig. S3.* Peak modelling and debris compensation analysis for the fluorescence data of the fresh preparation of *Oryza sativa* (A) and *Adenanthos sericeus* (B).

*Fig. S4.* Conventional histogram analysis for the fluorescence data of frozen preparations of *Oryza sativa* (P1) and *Adenanthos sericeus* (P2)

*Fig.* S5. Peak modelling and debris compensation analysis for the fluorescence data of the frozen preparation of *Oryza sativa* (A) and *Adenanthos sericeus* (B).

*Fig. S6.* Conventional histogram analysis for the fluorescence data of fresh preparations of *Oryza sativa* (P1) and *Hollandaea sayeriana* (P2).

*Fig. S7*. Peak modelling and debris compensation analysis for the fluorescence data of the fresh preparation of *Oryza sativa* (A) and *Hollandaea sayeriana* (B).

*Fig. S8.* Conventional histogram analysis for the fluorescence data of frozen preparations of *Oryza sativa* (P1) and *Hollandaea sayeriana* (P2).

*Fig. S9*. Peak modelling and debris compensation analysis for the fluorescence data of the frozen preparation of Oryza *sativa* (A) and *Hollandaea sayeriana* (B).

*Fig. S10.* Conventional histogram analysis for the fluorescence data of fresh preparations of *Oryza sativa* (P1) and *Macadamia tetraphylla* (P2).

*Fig. S11.* Peak modelling and debris compensation analysis for the fluorescence data of the fresh preparation of *Oryza sativa* (A) and *Macadamia tetraphylla* (B).

*Fig. S12.* Conventional histogram analysis for the fluorescence data of frozen preparations of *Oryza sativa* (P1) and *Macadamia tetraphylla* (P2).

*Fig. S13.* Peak modelling and debris compensation analysis for the fluorescence data of the frozen preparation of *Oryza sativa* (A) and *Macadamia tetraphylla* (B).

*Fig. S14.* Conventional histogram analysis for the fluorescence data of fresh preparations of *Oryza sativa* (P1) and *Macadamia jansenii* (P2).

*Fig. S15.* Peak modelling and debris compensation analysis for the fluorescence data of the fresh preparation of *Oryza sativa* (A) and *Macadamia jansenii* (B).

*Fig. S16.* Conventional histogram analysis for the fluorescence data of frozen preparations of *Oryza sativa* (P1) and *Macadamia jansenii* (P2).

*Fig. S17.* Peak modelling and debris compensation analysis for the fluorescence data of the frozen preparation of *Oryza sativa* (A) and *Macadamia jansenii* (B).

## Acknowledgements

The ARC Centre of Excellence for Plant Success in Nature and Agriculture provided funding (CE200100015) and Dr Shaun Walters technical assistance with access to the Flow cytometry facilities at the School of Molecular Bioscience, UQ. Dr Pauline Okemo and Dr Phoung Hoang from ARC Centre of Excellence for Plant Success in Agriculture and Nature grew the *Oryza sativa*.

## Authors’ contributions

AS: Conceptualisation, protocol testing, sample collection and processing, data collection, data analysis, drafting the manuscript, review and editing the manuscript, LS: Conceptualisation, sample collection, protocol testing, manuscript review and editing, AF: Conceptualisation, protocol testing and manuscript review and editing, RH: Conceptualisation, funding application, manuscript review and editing.

## Conflict of interest

The authors declare that they have no competing interests.

## Funding

AS was awarded a PhD Scholarship and RH was funded by the ARC Centre of Excellence for Plant Success in Agriculture and Nature (Grant CE200100015). LC was supported by a University of Queensland Retention Fellowship.

## Data availability

The dataset generated during the current study is available in the (“Genome size estimation of plants using frozen plant material”) flow repository database at (http://flowrepository.org/id/FR-FCM-Z6M4). The data supporting the conclusions of this article are included within the article and its additional files. Protocol was made available online at DOI: dx.doi.org/10.17504/protocols.io.e6nvwdq27lmk/v1 (Private link for reviewers: https://www.protocols.io/private/36483B59AE9611EE90E30A58A9FEAC02 to be removed before publication).

## Abbreviations

FCM: Flow cytometry
GS: genome size
CV: Coefficient of Variance
ANOVA: Analysis of variance
nls: Non-linear least square
RCS: Residual Chi-square
pg: picograms.

## Footnotes

1 Very fine grinding will damage the intactness of the nuclei, whereas too coarse will yield fewer nuclei. For larger genomes, keeping the powder coarsely ground without any chunks or small leaf pieces is recommended. After grinding, secure the plant material at or below −80°C until processed further.

## Notes

### Competing Interest Statement

The authors have declared no competing interest.

